# Nonparametric analysis of contributions to variance in genomics and epigenomics data

**DOI:** 10.1101/314112

**Authors:** David M. Moskowitz, William J. Greenleaf

**Affiliations:** Biomedical Informatics Training Program, Stanford University School of Medicine, Stanford, CA 94305, USA; Department of Genetics, Stanford University School of Medicine, Stanford, CA 94305, USA

## Abstract

Functional genomics studies, despite increasingly varied assay types and complex experimental designs, are typically analyzed by methods that are unable to identify confounding effects and that incorporate parametric assumptions particular to gene expression data. We present MAVRIC, a nonparametric method to quantify variance explained by experimental covariates and perform differential analysis on arbitrary data types. We demonstrate that MAVRIC can accurately associate covariates with underlying data variance, deliver sensitive and specific identification of genomic loci with differential counts, and provide effective noise reduction of large-scale consortium data sets.

## Main text

Rapid advances in sequencing technology have enabled the development of assays interrogating chromatin structure^1^, DNA methylation^2^, and DNA/protein interactions^3^, with data increasingly generated by large, multi-institutional projects^4^. However, analysis pipelines often hew to the paradigms established by early microarray and RNA-seq methods, such as limma^5^ and DESeq^6^. These workflows involve two main steps: First, quality control analysis, often through application of principal component analysis (PCA) to verify that variance between samples aligns primarily with the biology of interest. Second, analysis of differential expression, quantified using a linear model to individually test genes for nonzero fold-changes in expression across samples. Formulating these linear models requires users to explicitly encode every term they expect to affect expression, and any interactions thereof. Because of their rigid definitions, the linear models can yield misleading results when the appropriate technical covariates are omitted, and they will generally fail to provide insight into confounding effects^7^.

Sequencing sample and batch heterogeneity, particularly in large research projects, can induce complex technical artifacts^8^, posing a challenge in differential analysis. To correct for technical effects more broadly, researchers must employ independent methods that use factor analysis to model latent variables^9-11^. However, such approaches can inadvertently capture and blunt estimates of real biological effects^12^. Moreover, the magnitude and impact of the correction is dependent on the number of factors in the model^12^, which the user is required to specify.

Additionally, most differential expression analysis tools are poorly suited to the broad range of modern sequencing assays, since those methods typically incorporate parametric assumptions particular to gene expression data. They thus implicitly require that individual loci be of fixed sequence composition and length^13^. Such assertions are inapplicable to epigenetic data, as the precise boundaries of epigenomic features can vary across samples^14^.

To provide an integrative, scalable method for identifying technical confounding and performing assay-agnostic differential analysis, we have developed MAVRIC (Measuring Association between VaRIance and Covariates). MAVRIC is a nonparametric statistical method that quantifies signal-to-noise, identifies statistical confounding between experimental covariates, and performs pairwise differential analysis. Because MAVRIC does not rely on assumptions about count distributions or model structure, the algorithm offers a framework suitable to diverse assay types and expansive experimental designs with numerous covariates. MAVRIC extends the use of PCA in quality control analysis, aiming to make PCA interpretable and actionable by associating PCs with experimental covariates and defining PC-based “axes of variance” for differential analyses. Crucially, in contrast to many other analysis methodologies, MAVRIC focuses on quantifying and visualizing the degree to which a data set successfully captures the impact of the biological feature being assayed, rather than artifactual or stochastic differences.

As input, MAVRIC takes a *P* by *N* data matrix and an *N* by *M* design matrix, for *N* samples, *P* features (e.g. genes or loci), and *M* known experimental covariates, where each covariate is categorical (Fig. 1a) or continuous (see Methods). The data matrix need not contain raw counts. First, MAVRIC performs PCA on the data matrix, after (optionally) using a variance-stabilizing transformation to correct for the heteroskedasticity of sequencing data^13^. Unlike common RNA-seq analysis tools, MAVRIC is also compatible with arbitrary normalization methods. Next, a built-in gene selection workflow can be used to retain only features with high variance in counts. Then, MAVRIC iterates over the covariates in the design matrix, greedily associating each covariate with the maximal set of PCs that spatially segregate covariate categories more than expected by chance (see Methods, Supplementary Fig. 1a,b).

**Figure 1:**
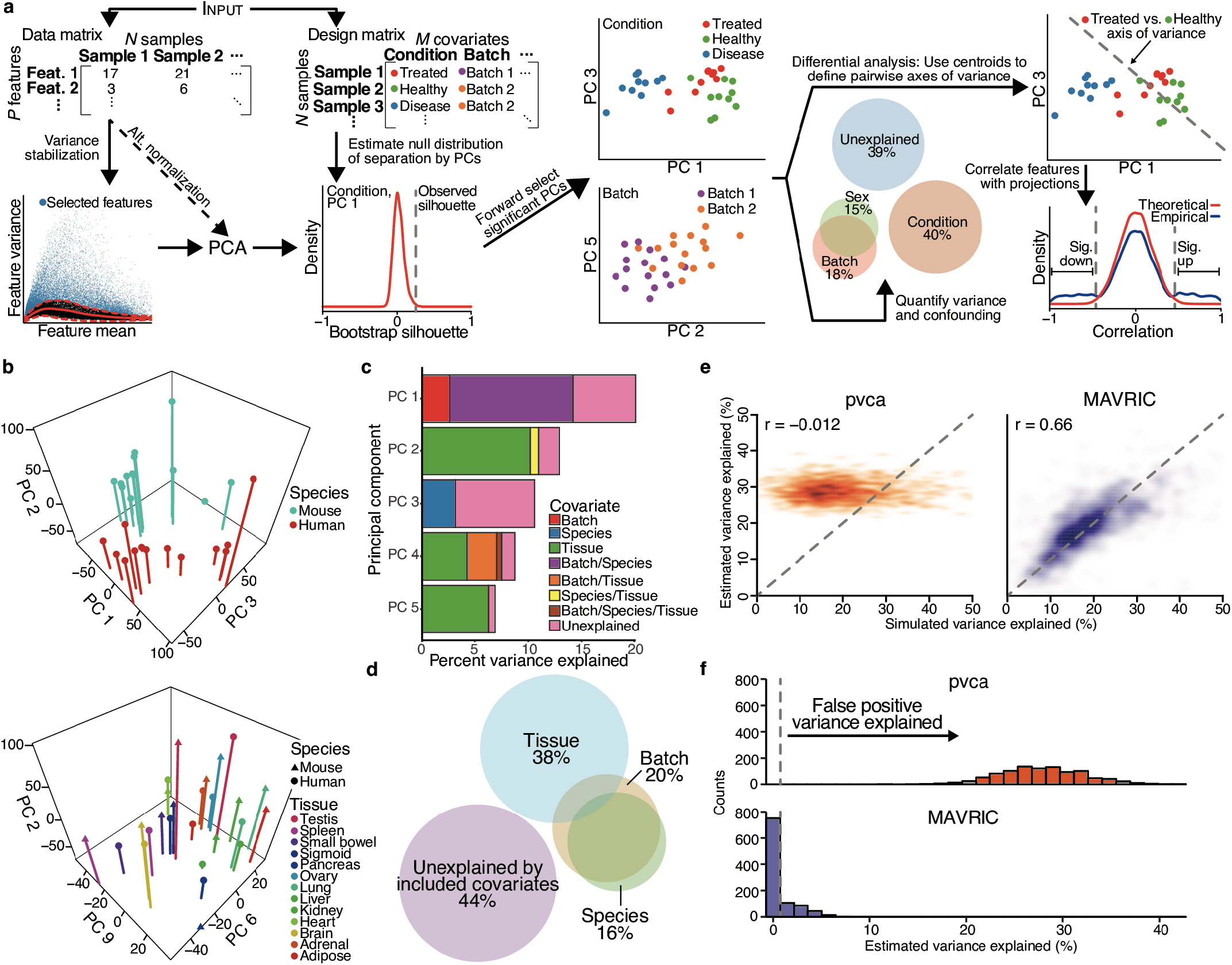
Associating PCs with covariates to quantify sources of variance. (a) The MAVRIC algorithm: (1) input a data matrix and a covariate matrix, (2) optionally normalize data and select high-variance features, (3) perform PCA, (4) associate PCs with covariates to quantify contributions to variance, and (5) define “axes of variance” for differential analysis. (b) Top three PCs identified by MAVRIC as associated with species-specific and tissue-specific effects in previously published RNA-seq data^15^. Next three PCs depicted in Supplementary Fig. 1d. (c) Quantification of variance explained by known covariates across first five PCs of RNA-seq data from (b). (d) Total variance explained by known covariates, and residual unexplained variance, as aggregated across all PCs of RNA-seq data from (b) and (c). Overlaps in Euler diagram represent statistical confounding between covariates. (e) Accuracy of pvca’s (left) and MAVRIC’s (right) estimation of variance explained by a covariate in 1,000 simulated RNA-seq experiments. (f) Estimates of variance explained by extraneous covariates, as assessed by pvca (top) versus MAVRIC (bottom), across 1,000 simulations of RNA-seq data where all variance is stochastic.

As output, MAVRIC quantifies the variance that the covariates explain, and how those effects are confounded, based on the PC associations. Whereas typical workflows use PCA only to visualize the biology of interest in the context of the first two PCs, MAVRIC provides a more complete picture of the underlying sources of variance and their relationships to lower-rank PCs. As a demonstration of this functionality, we re-analyzed RNA-seq data generated across multiple tissues from mouse and human^15^. Previous analysis of this data set indicated that differences between species, rather than tissues, drove higher-rank PCs, but concerns about confounding between species and batch effects were subsequently raised^16^. The MAVRIC results support species-specific effects as driving high-rank PCs, with tissue-specific differences instead associated with lower-rank PCs (Fig. 1b, Supplementary Fig. 1d). Additionally, MAVRIC identifies statistical confounding between batch and species effects (Figs. 1c,d). Finally, MAVRIC quantifies the relative contributions to count variance of the species, batch, and tissue effects, along with the variability that is not significantly attributable to any of those covariates. The residual noise could represent stochastic expression differences, variability between tissue donors, or other technical effects (“unexplained,” Figs. 1c,d). Thus, MAVRIC’s quantification of variance and confounding, and its corresponding Euler diagram output, offers an interpretable analysis that allows researchers to assess whether an experiment has effectively captured the biology of interest, and if there are potential technical issues.

Unlike highly structured linear models, MAVRIC considers each covariate independently. Consequently, adding terms to the design matrix only affects estimates of confounding, without perturbing the analysis results of other covariates. Using simulated data, we found that the estimates of variance explained provided by MAVRIC were accurate and insensitive to the inclusion of superfluous covariates (Figs. 1e,f). We compared these results to those of pvca^17^, another method for estimating contributions to variance, based on linear models. We observed that pvca consistently demonstrated lower accuracy, and often attributed variance to modeled terms even when they did not actually affect the simulated count variance (Figs. 1e,f).

MAVRIC further performs differential analyses by defining axes of variance for pairs of categories within a covariate. For a pairwise comparison, the axis of variance is the line defined by the centroids of the two categories in the subspace of PCs that MAVRIC associated with the covariate. Onto this axis, MAVRIC projects the points of all the samples across the entire data set, and those projected values are correlated with the original values from the data matrix (Supplementary Fig. 1c; see Methods). Every sample type in the analysis contributes to the basis set defining the subspace that best separates the covariate’s categories, and likewise all samples are expected to capture information relevant to the biological variation of interest. Therefore, MAVRIC uses all the samples in the data set when performing pairwise comparisons. While this design does not affect estimated differential effect sizes, it can result in greater sensitivity when identifying features correlated with the axis of variance (see Methods). A significant correlation between feature values and projected values indicates that the feature variance aligns with the patterns detected by PCA on the full data set. Such features are called as differential elements for that pairwise comparison. In this sense, unlike traditional RNA-seq methods that calculate parameters and perform statistical tests per gene, MAVRIC’s results statistically leverage modules of features with coherent changes, since those modules determine the directions of the PC vectors.

We evaluated MAVRIC by assessing its differential analyses in comparison to a widely used RNA-seq method, DESeq2^13^. On simulated data generated using the parametric assumptions of DESeq2, we determined that MAVRIC’s performance was competitive across a broad range of simulated expression changes, while also finding an order-of-magnitude fewer false positives when no differences were present (Figs. 2a,b, Supplementary Fig. 2a). We then tested both methods on real, previously published ATAC-seq data^18^, which were not expected to conform to the parametric assumptions used for RNA-seq data. The ATAC-seq data cataloged chromatin accessibility in naive, central memory, and effector memory human CD8 T cells, and we applied both MAVRIC and DESeq2 to identify loci more accessible in the effector memory than the central memory cells. We found that MAVRIC recapitulated the majority of the significant results from DESeq2, while identifying substantially more changes overall (Fig. 2c). We next calculated sequence enrichment for transcription factor binding motifs in each of three sets of loci identified as differentially accessible: unique to MAVRIC (purple circle, Fig. 2c); unique to DESeq2 (green circle, Fig. 2c); and overlapping (brown circle, Fig. 2c), which we considered a gold standard (Supplementary Fig. 2b). We compared the gold-standard motif enrichment p-values to those of the two other sets and observed that motif enrichment p-values for MAVRIC exhibited a significantly higher area under the receiver operating characteristic curve (AUC) than did the p-values for DESeq2 (Fig. 2d). Gene ontology enrichment of the differentially accessible peaks also showed high concordance between the results for the overlapping peaks and the peaks unique to MAVRIC, while the peaks unique to DESeq2 produced no statistically significant categories (Supplementary Fig. 2c).

**Figure 2:**
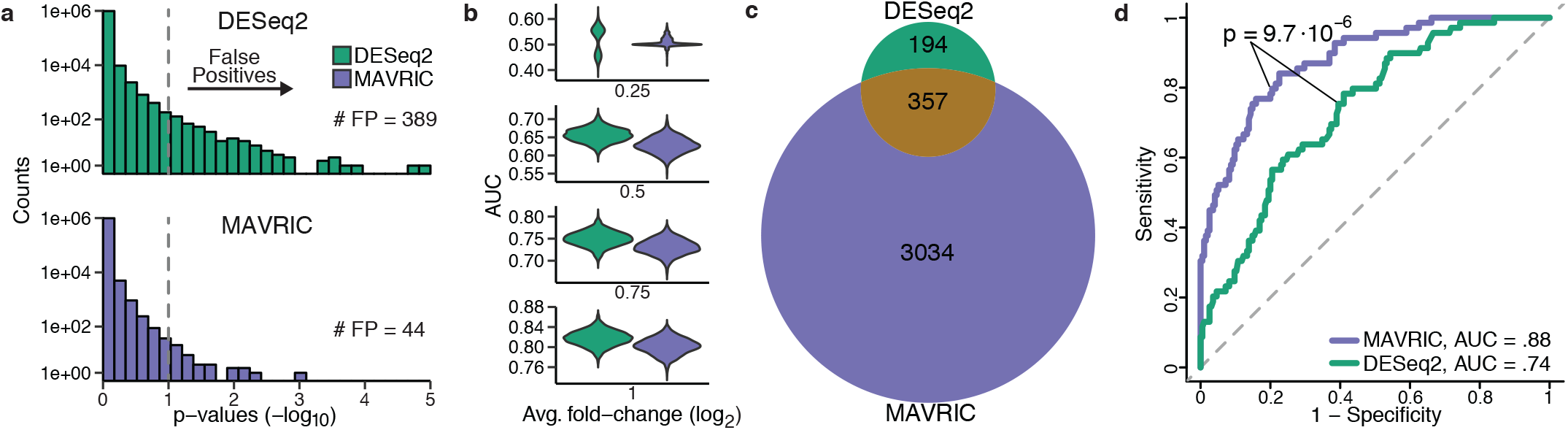
Evaluation of differential analysis results on simulated and real data. (a) False positives from DESeq2 (top) and MAVRIC (bottom), across 1,000 simulated RNA-seq experiments, where no genes are differentially expressed. Dashed, vertical line corresponds to an adjusted p-value of 0.1. (b) Performance of DESeq2 (green) versus MAVRIC (purple) in identifying differentially expressed genes, across 1,000 simulated RNA-seq experiments, with counts drawn from negative binomial distributions. Y-axes give distributions of AUCs, and each pane corresponds to a different average simulated fold-change. (c) Venn diagram of ATAC-seq peaks found to be more accessible in human effector memory CD8 T cells (EM), as compared to central memory CD8 T cells (CM), as assessed by applying MAVRIC and DESeq2 to previously published data^18^. (d) Comparison of areas under the receiver operating characteristic curves (AUCs) in recovering statistically significant transcription factor binding motif enrichment from gold-standard ATAC-seq peaks more accessible in EM than CM (357 peaks in brown overlap from (c)). Purple curve corresponds to enrichment p-values in MAVRIC-specific peaks (3,034 peaks in purple in (c)) and green curve corresponds to enrichment p-values in DESeq2-specific peaks (194 peaks in green in (c)). P-value between AUCs is from DeLong’s test.

Next, we applied MAVRIC to large-scale consortium data, to test its effectiveness at distinguishing biological signal from the noise of experimental heterogeneity of data generated by multiple investigators at different institutions. Using RNA-seq data of 29 human tissues from the GTEx project, which aims to connect tissue-specific gene expression levels to expression quantitative trait loci (eQTLs)^19^, we found that correlating samples based on raw counts and applying *k*-means partially separated the samples by tissue (Fig. 3a). Repeating this analysis on counts adjusted by sva^11, 20^, to correct for latent factors affecting the expression measurements and identify differentially expressed genes, we observed that the clustering more accurately reflected tissue types, while reducing the total data variance by 9.6% (Fig. 3b). When we instead applied MAVRIC to the counts, to select for high-variance features and PCs associated with the tissue covariate, we determined that the clustering improved more than with sva, while reducing variance by 10.4% relative to the raw data (Fig. 3c). The high-variance genes identified by MAVRIC were also more likely to be differentially affected by eQTLs across tissues than were the top differentially expressed genes identified by sva’s workflow (see Methods, Supplementary fig. 3). Thus, compared to sva, MAVRIC offered a data correction that provided superior clarity of the biological differences of interest, while retaining a similar fraction of total variance.

**Figure 3:**
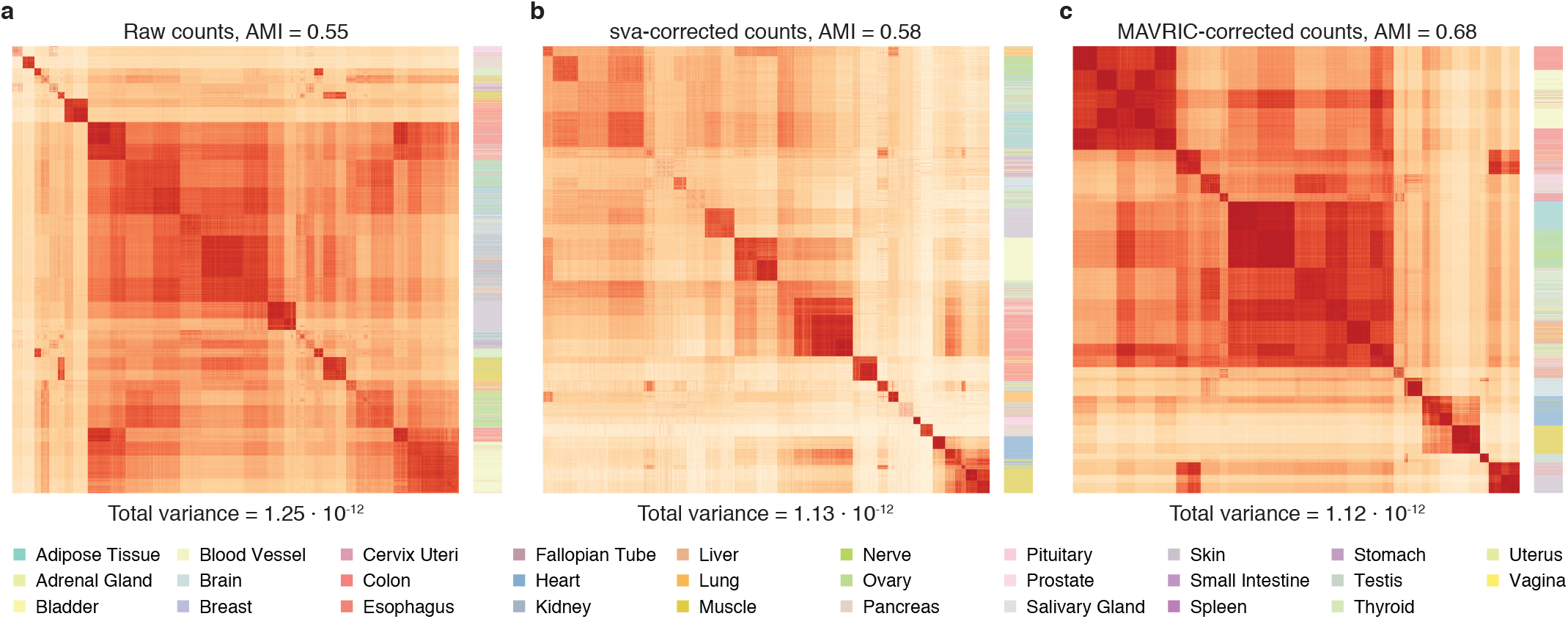
Noise reduction in large-scale consortium data. (a) Correlation matrix of RNA-seq samples from the GTEx project^19^, with samples ordered by k-means clustering. Color bar on right represents tissues. AMI refers to adjusted mutual information, which measures agreement between tissue annotations and cluster assignments, where 1 represents perfect agreement and 0 represents the agreement expected by chance. (b) As in (a), for the top differentially expressed genes identified after correction with sva. (c) As in (a), for genes identified by MAVRIC as having high variance in expression, and counts corrected by dropping PCs not associated with tissue-driven effects. The number of genes used is equal to that in (b).

By associating PCs with experimental covariates, MAVRIC can identify sources of variance, perform differential analysis, and reduce artifactual variation, without requiring that the user make explicit assumptions about which covariates are impactful. By focusing on data visualization, MAVRIC’s outputs maximize interpretability, providing users with a comprehensive overview of the biological and technical effects captured by functional genomics experiments. We expect that MAVRIC will provide a valuable complement to gene expression-oriented methods, particularly for use with newer assay types and in more expansive experimental designs.

---

**Supplementary Figure 1:**
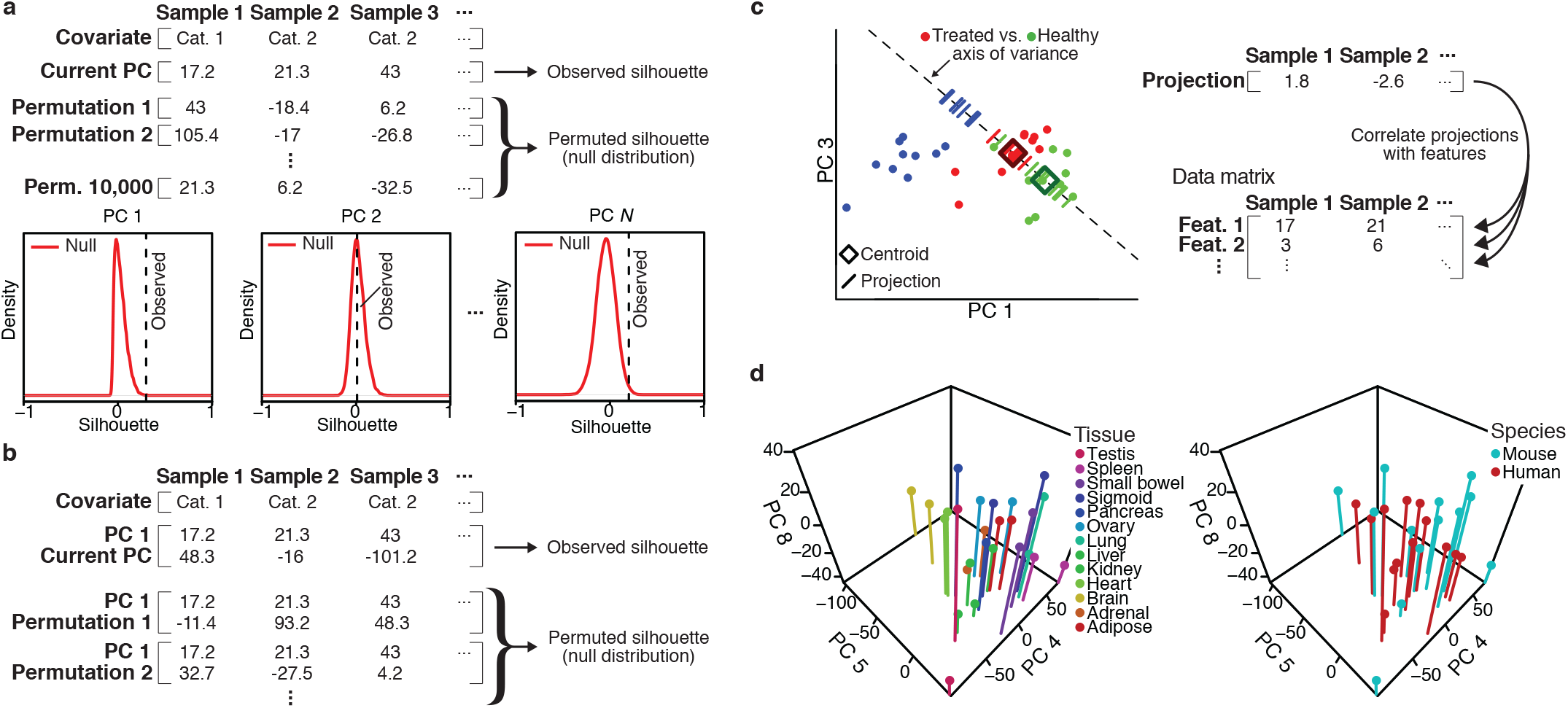
Connecting variance to experimental covariates. (a) To statistically quantify the association between a PC and a covariate, MAVRIC uses the average silhouette of the points with respect to the category labels. The silhouette measures how segregated the categories are within the PC’s subspace (see Methods). MAVRIC then obtains a p-value for the observed average silhouette by permuting the coordinates 10,000 times and recalculating the average silhouette, forming a null distribution. The p-value of the observed average silhouette with respect to the null distribution tests the hypothesis that the PC offers a coordinate space in which the categories are segregated more than expected by chance. When more than one PC exhibits a significant p-value (based on a user-specified alpha), MAVRIC selects the single PC for which the difference between the observed average silhouette and the expected average silhouette is maximal, where the expected average silhouette is the mean of the null distribution. (b) PCs beyond the first are added to a covariate’s association subspace by forward selection. In particular, during the permutations, only the newly added PC’s coordinates are permuted, while those that MAVRIC has already associated with the covariate remain constant. Thus, the p-value tests the hypothesis that adding another PC to the current subspace improves the segregation between the categories more than expected by chance. (c) MAVRIC performs differential analysis by defining an “axis of variance” for a particular pairwise comparison between categories. The axis of variance is the line given by the two categories’ respective centroids in the subspace of associated PCs. Each sample’s coordinate in the associated PC subspace is projected onto that axis, yielding a univariate point for every sample in the data set. These projected values are then correlated with the values from the original data matrix. A high absolute correlation indicates that the variance for a feature aligns with the variance between the two categories, and thus that that feature is differentially expressed. (d) Fourth-through sixth-ranked PCs identified by MAVRIC as associated with tissue-specific effects in previously published RNA-seq data^15^.

**Supplementary Figure 2:**
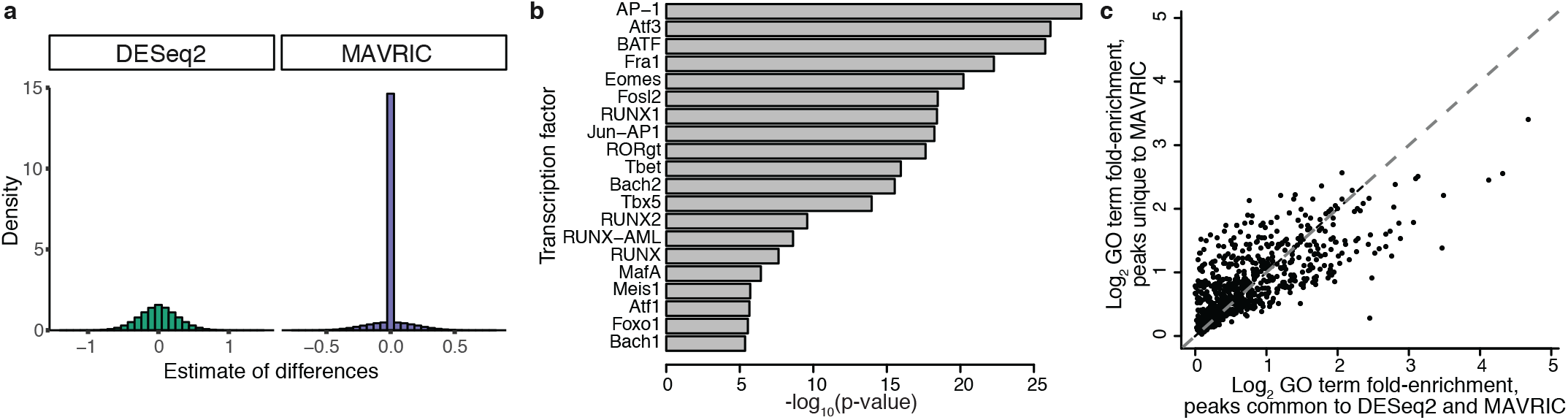
Comparison of MAVRIC and DESeq2 on ATAC-seq data. (a) Distributions of estimates of effect sizes of differential expression, in simulated data without any differential expression (see Methods), for DESeq2 (left) and MAVRIC (right). For DESeq2, the x-axis is on the scale of log2 fold-changes. For MAVRIC, the x-axis is on the scale of correlations. (b) Transcription factor (TF) binding motif enrichment, for the top enriched TF motifs in the ATAC-seq peaks identified by both DESeq2 and MAVRIC as more accessible in human CD8 effector memory T cells than central memory cells (brown circle, Fig. 2c). The results are concordant with those of the original study^18^, and were thus used as a gold standard. (c) Gene ontology (GO) enrichment for peaks identified as differentially accessible by both MAVRIC and DESeq2 (x-axis; brown circle, Fig. 2c), versus GO enrichment for peaks identified as differentially accessible only by MAVRIC (y-axis; purple circle, Fig. 2c). Each point represents a GO category that was significantly enriched in at least one of the two peak sets at an alpha of .05 after multiple hypothesis correction. Pearson correlation = 0.70. The peaks identified as differentially accessible only by DESeq2 did not produce any significantly enriched GO categories.

**Supplementary Figure 3:**
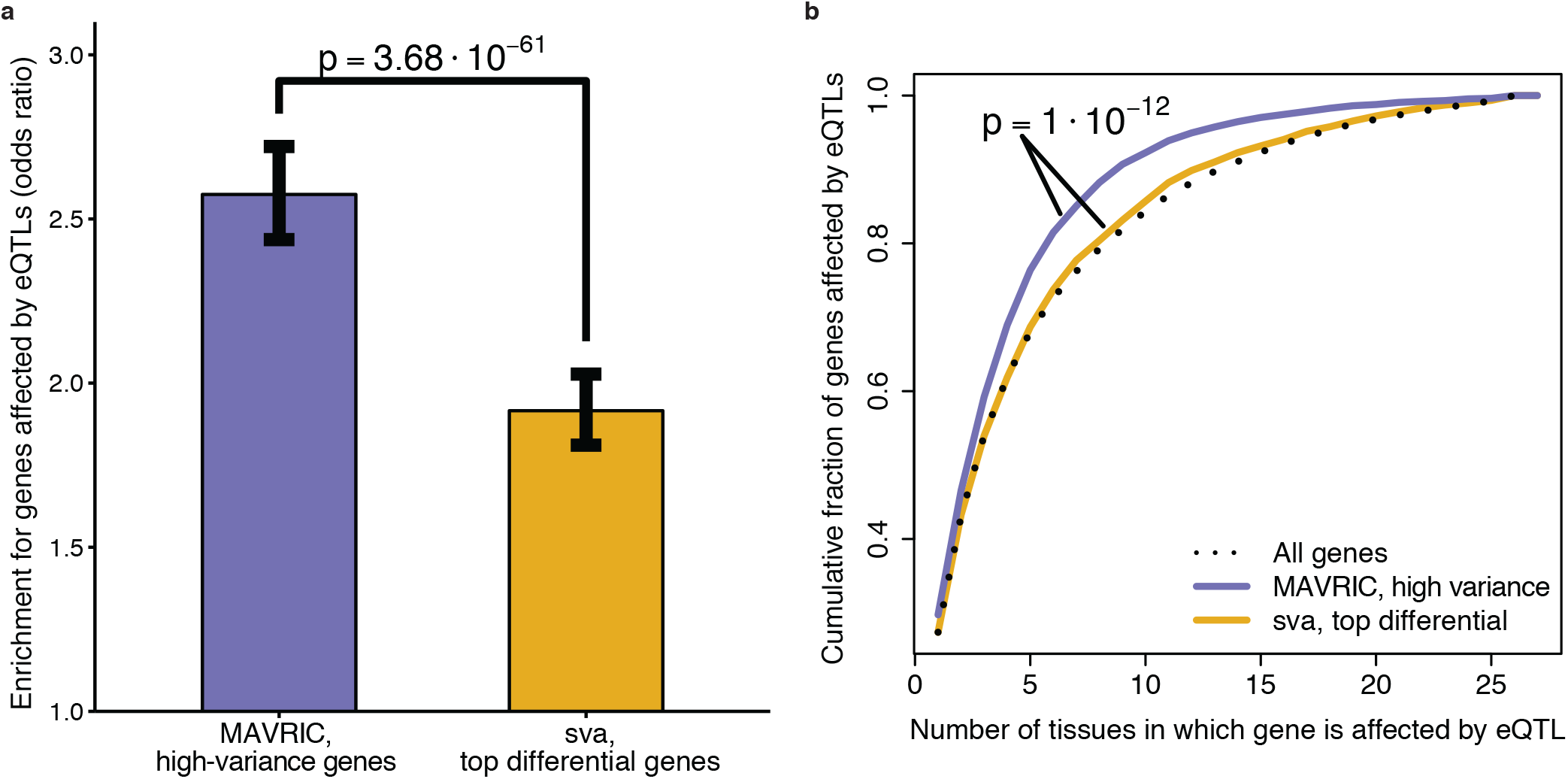
Representation of genes affected by eQTLs following correction of GTEx data by sva versus by MAVRIC. (a) Odds ratios for likelihood that a selected gene had expression affected by an eQTL, for MAVRIC (left) and sva (right), as compared to all genes in the GTEx data set^19^. For MAVRIC, selected genes are those identified by its feature selection workflow as having high variance (see Methods). For sva, selected genes are an equal number of the top genes (as measured by smallest p-value) with significantly differential expression across tissues following data correction (see Methods). (b) Distributions of numbers of tissues in which gene expression is affected by an eQTL. Sets of genes used for MAVRIC (purple) and sva (gold) are as in (a). Dotted black line gives distribution across all genes in the GTEx data.

## Methods

### Inputs to MAVRIC

MAVRIC has two required inputs, a data matrix and a design matrix (Fig. 1a). The data matrix is of dimension *P* × *N*, for *P* features measured across *N* samples. For RNA-seq, the features would represent genes; for ChIP-seq or ATAC-seq, the features would represent peaks. The design matrix is of dimension *N* × *M*, for *M* covariates annotated across the same set of samples. The covariates reflect biological and technical attributes of the sample (e.g. treatment, sex, RIN), and can represent either categorial or continuous variables. MAVRIC requires that each categorical covariate has at least two categories, and that each category has at least two samples.

### Data preprocessing in MAVRIC

The data matrix is optionally initially processed with a variance-stabilizing transformation to reduce heteroskedasticity, thereby likewise reducing the extent to which features with high means act as the primary drivers of the directions of the PC vectors. To avoid any parametric assumptions, the logarithm can be applied, or the user can employ a normalization with parametric assumptions particular to RNA-seq. Next, MAVRIC attempts to improve statistical power by eliminating features with low variance, which are unlikely to be differential. For the feature selection process, MAVRIC fits, to the mean versus variance trend, an iteratively reweighted loess function, with weights given by a redescending M-estimator (Tukey’s biweight) based on the residuals. For point *i* with residual *r(i)*, the weight *w(i)* is:

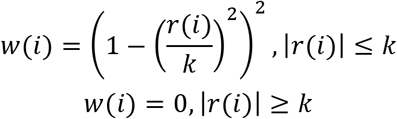

The constant *k* controls robustness and here is set to the typical value of 4.685, which provides desirable asymptotic efficiency. Thus, features with high residuals are assigned lower weights, yielding a fit where high-variance features have low statistical leverage. Features with variance above the 95% confidence interval of the loess fit are considered to have higher than expected variance given their means; the remaining features are discarded.

The normalization and feature selection steps described above are both optional and flexible: The user is also able to normalize the data prior to inputting it to MAVRIC, and may further decide whether or not to employ a variance-stabilizing transformation. For feature selection, the user has the option of specifying that MAVRIC retain the *v* features with the highest variance (after normalization), rather than using the loess fit to control for any residual mean/variance dependence.

### Using MAVRIC to associate PCs with experimental covariates

On the normalized data matrix, MAVRIC performs PCA, and then iterates over the covariates in the design matrix, associating each covariate with a subset of the PCs. Certain PCs are discarded at the outset, according to a user-specified eigenvalue threshold: Any PC that explains less variance than the threshold is discarded without evaluating its associations with the covariates.

For categorical covariates, MAVRIC uses forward selection to establish the maximal set of PCs for which the spatial separation between the categories is greater than expected by chance. The use of forward selection helps reduce average run-time, particularly for large-scale consortium data with many samples, because the majority of PCs are generally expected to correspond to inter-sample differences and stochastic variability, with only a small number of PCs attributable to variance associated with each covariate. In the forward selection workflow, for each PC, the silhouette statistic^21^, averaged across all samples, is used to quantify the separation between the categories. For a sample *i* in category *A*, the silhouette, *s(i)*, assesses how close *i* is to the others in its category, relative to the samples in the spatially nearest category:

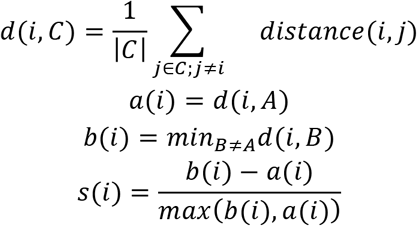

The function *distance(i,j)* refers to the Euclidean distance between points *i* and *j* in the current PC subspace. The observed average silhouette is compared against the null distribution generated by average silhouette statistics calculated on 10,000 permutations of the PC coordinates. If there is a PC for which the one-sided p-value for the observed average silhouette is significant, that PC is considered to be associated with the covariate; if there is more than one such PC, only the single PC with the greatest difference between the observed average silhouette and the expected average silhouette is taken. The process then repeats on the remaining PCs, with null distributions calculated by permuting only the newly added coordinates, to test whether adding that dimension to the current PC subspace improves the separation more than expected by chance, given the current subspace. When there are no further PCs that can be added to improve the separation, the process ends, and the next covariate is considered.

For continuous covariates, significant associations are established using the lasso. The lasso is not likewise used with categorical covariates for compatibility with the small sample sizes characteristic of sequencing experiments. In the lasso, For responses *y* and PC coordinates *X*, the regression coefficients, β, are given by:

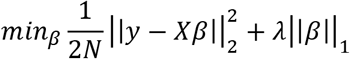

The regularization parameter, λ, is determined by cross-validation. The number of folds in the cross-validation is set such that there are at least three samples in each fold, up to a maximum of 10 folds. When fewer than 15 samples are supplied, leave-one-out cross-validation is employed.

### Estimating covariates’ contributions to data variance with MAVRIC

Using the associations between PCs and covariates, MAVRIC calculates each covariate’s contribution to the total data variance. To determine contributions to variance, MAVRIC calculates a weighted sum of eigenvalues for each covariate, where the eigenvalues are those that correspond to the PC vectors. For PCs associated with the covariate, the weight is the within-group sum-of-squares (WGSS) divided by the total sum-of-squares (TSS), within that PC coordinate’s subspace; for PCs not associated with the covariate, the weight is 0. The contribution to variance *V(A)* for covariate *A* with categories *C* is as follows:

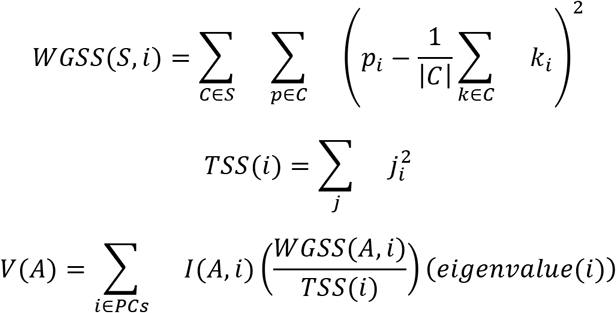

*I(A,i)* is an indicator variable that takes value 1 if PC *i* is associated with covariate *A*, and 0, otherwise. The function *eigenvalue(i)* returns the variance explained by PC *i*. Confounding between covariates is given by the variance shared across overlapping associated PCs:

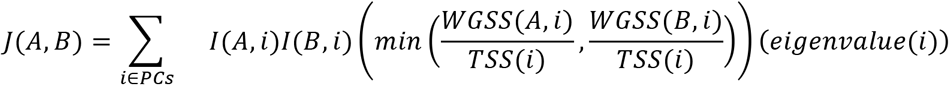

Residual variance is considered to be unexplained by the supplied covariates:

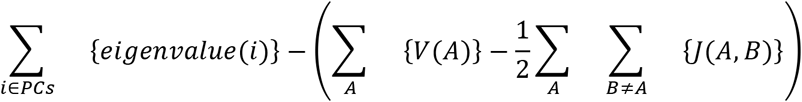

### Performing differential analysis with MAVRIC

MAVRIC leverages associations between covariates and PCs to perform pairwise differential analysis between categories within a covariate. Having identified a PC subspace providing separation between the categories, MAVRIC seeks to identify features for which the variance in counts across samples matches the locations of the samples within that subspace. To achieve this goal, MAVRIC first reduces the spatial position of each sample within the PC subspace to a univariate value. That value corresponds to whether the sample is more similar to the average of one category versus the other. MAVRIC then correlates those values to the original data, for each feature in the data matrix, attaining estimates of whether the original count variance for a feature aligns with the spatial distances between samples within the PC subspace associated with the covariate.

Specifically, to evaluate pairwise differential expression, MAVRIC begins by defining an “axis of variance” for each pair of categories, for each covariate. For a given pair of categories, the axis of variance is the line given by the pairs’ respective centroids in the subspace of PCs associated with the covariate. The zero point of the line is defined to be the midpoint between the centroids. Onto that line, MAVRIC projects the each sample’s point, generating a univariate value for each sample. This procedure is carried out for every sample, including those that are not members of the categories in the pairwise comparison. This design leverages the full data set when identifying elements that vary along the axis of variance, thus increasing statistical power (without affecting estimates of effect sizes; see below). The projection of point *p* onto the line defined by centroids *c* and *d*, with midpoint *m*, is given by:

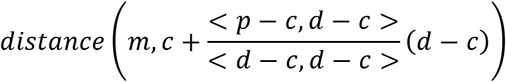

The unsigned distances are converted to signed values based on to which of the two centroids the projected point is closer. The signed values are then correlated with each feature from the original data matrix. Features for which that correlation is statistically significant are considered to be differential for that pairwise comparison.

Effect sizes are determined as typical fold-changes, except that, rather than using the original data matrix, they are calculated based on a reconstructed data matrix retaining only those PCs that MAVRIC identified as being associated with the covariate. By dropping out irrelevant PCs, MAVRIC estimates effect sizes that are natively corrected for unwanted variation.

### Analysis of ENCODE RNA-seq data

ENCODE RNA-seq data across 13 tissues from mouse and human, covering 14,744 genes, were drawn from a previously published study^15^. The counts were translated to RPKMs and log-transformed, after which they were inputted to MAVRIC. To facilitate a direct comparison between MAVRIC’s results and the previously published analysis, MAVRIC’s feature selection was not used.

### Data simulations

Simulations were generated in sets of 1,000, across 27 samples and 1,000 features, annotated with 3 covariates, each with 3 categories. For all simulations, the framework was derived from the DESeq2 function makeExampleDESeqDataSet, which simulates counts from a negative binomial distribution, where each gene has a randomly generated mean and fold-change, with overdispersion as a function of the mean. This function was generalized to allow for fold-changes across multiple categories and covariates, and Gaussian noise was added both to the final counts, as well as per sample to the means and overdispersion parameters, such that each sample was generated from a negative binomial distribution with slightly different parametrization. The Gaussian noise parameters were chosen such that, after adding noise, the perturbation was +/- 10% of the original value, on average. For feature *j* of sample *i*, in a category *v* with average log_2_ fold-change *k* relative to the baseline simulated expression level, the counts *c* are distributed as:

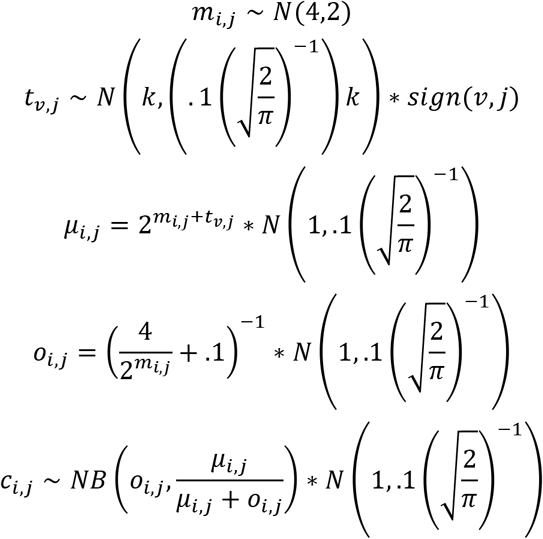

The function *sign(v,j)* controls whether the feature is down- or up-regulated relative to the baseline mean, and takes on values 1 and −1 with equal probability. Because the *c_i,j_* are intended to represent count data, the final values are rounded to the nearest integer.

When benchmarking accuracy relative to pvca, counts were first simulated with all fold-changes between categories set to zero. Then, for each simulation, 333 genes were randomly selected and had their counts redrawn such that, with respect to one of the covariates, all three categories were of different mean. The new counts were then scaled such that the new overall mean across all samples equalled the mean of the original parametrization. For both MAVRIC and pvca, the values plotted for estimates of variance explained were with respect to the covariate for which the categories’ means had nonzero fold-changes.

When comparing the baseline false positives rates of MAVRIC to those of pvca and DESeq2, simulations were generated with nonzero fold-changes across all categories for every covariate. Estimates of variance explained were summed across all covariates. False positive rates for differential expression were with respect to a single pairwise comparison for one covariate.

AUCs for differential expression were calculated across four sets of simulations, each with a different average pairwise fold-change across categories from a single covariate. For each gene in these simulations, a category was selected with .2 probability to have a mean different from the baseline mean. The AUCs plotted were with respect to a single pairwise comparison within the covariate for which the categories’ means had nonzero fold-changes.

### Analysis of ATAC-seq data

Previously published ATAC-seq data from human CD8 T cell subsets^18^ were reanalyzed using MAVRIC. These results were compared to those of DESeq2, the method used for differential analysis in the original study. The pairwise comparison for which the results were compared was for ATAC-seq peaks more accessible in effector memory (EM) cells than central memory (CM) cells; this pairwise comparison was chosen because DESeq2 was unable to discover many significant changes in the published results. Here, changes were considered to be statistically significant based on an alpha of .01, after multiple hypothesis correction with the Benjamini-Hochberg procedure.

Transcription factor (TF) binding motif enrichment was assessed with HOMER^22^, using non-significant peaks as the background set. True positives for the AUCs were considered to be those motifs that were significantly enriched in the peak set identified as differentially accessible by both MAVRIC and DESeq2.

Gene ontology (GO) analysis was performed using GREAT^23^. Categories were considered to be significantly enriched based on an alpha of .05.

### Analysis of GTEx RNA-seq data

RNA-seq data and annotations of tissue-specific eQTLs from version 6 of the GTEx project were downloaded via the GTEx portal^19^. To fulfill the requirements for running MAVRIC, samples were discarded if they were members of categories for which there were fewer than three samples. This preprocessing yielded a total of 6,152 samples.

When running MAVRIC, the counts were first log-transformed, with a pseudocount of 1 added. MAVRIC’s feature selection workflow retained 7,594 genes with greater variance than expected by chance, given their respective means. To calculate the correlation matrix, PCs were discarded if they were not found to be associated with differences across tissues, and only those genes retained by feature selection were used.

When running sva, the method designed for sequencing data (svaseq) was used, parametrized to preserve variance across tissues. An F-test was used, in accordance with sva’s user guide, to quantify the likelihood that a gene was differentially expressed across some pair of tissues. To enable a direct comparison with MAVRIC, only the top 7,594 genes (as defined by smallest p-value) were used to calculate the correlation matrix.

Samples were clustered based on the correlation matrix using *k*-means clustering, with *k* set to 29. The fixed cluster number was chosen to be equal to the number of tissues represented in the data set. Adjusted mutual information was calculated for clusters with respect to the tissue annotations. This measure was chosen based on prior work demonstrating that it is the preferred statistic for comparing clusterings when there are unbalanced and small class sizes^24^, as is the case for the tissue annotations used.

